# Transient Polycomb activity represses developmental genes in growing oocytes

**DOI:** 10.1101/2022.09.18.508436

**Authors:** Ellen G. Jarred, Zhipeng Qu, Tesha Tsai, Ruby Oberin, Sigrid Petautschnig, Heidi Bildsoe, Stephen Pederson, Qing-hua Zhang, Jessica M. Stringer, John Carroll, David K. Gardner, Maarten van den Buuse, Natalie A. Sims, William T. Gibson, David L. Adelson, Patrick S. Western

## Abstract

**Background:** Non-genetic disease inheritance and offspring phenotype is substantially influenced by germline epigenetic programming, including genomic imprinting. Loss of Polycomb Repressive Complex 2 (PRC2) function in oocytes causes non-genetically inherited effects on offspring, including embryonic growth restriction followed by post-natal offspring overgrowth. While PRC2 dependent non-canonical imprinting is likely to contribute, less is known about germline epigenetic programming of non-imprinted genes during oocyte growth. In addition, *de novo* germline mutations in genes encoding PRC2 lead to overgrowth syndromes in human patients, but the extent to which PRC2 activity is conserved in human oocytes is poorly understood.

**Results:** In this study we identify a discrete period of early oocyte growth during which PRC2 is expressed in mouse growing oocytes. Deletion of *Eed* during this window led to the de-repression of 343 genes. A high proportion of these were developmental regulators, and the vast majority were not imprinted genes. Many of the de-repressed genes were also marked by the PRC2-dependent epigenetic modification histone 3 lysine 27 trimethylation (H3K27me3) in primary-secondary mouse oocytes, at a time concurrent with PRC2 expression. In addition, we found H3K27me3 was also enriched on many of these genes by the germinal vesicle (GV) stage in human oocytes, strongly indicating that this PRC2 function is conserved in the human germline. However, while the 343 genes were de-repressed in mouse oocytes lacking EED, they were not de-repressed in pre-implantation embryos and lost H3K27me3 during pre-implantation development. This implies that H3K27me3 is a transient feature that represses a wide range of genes in oocytes.

**Conclusions:** Together, these data indicate that EED has spatially and temporally distinct functions in the female germline to repress a wide range of developmentally important genes, and that this activity is conserved in the mouse and human germlines.

## Background

Epigenetic modifications, including DNA methylation and histone modifications, regulate chromatin packaging and underlie long-term cell-specific gene transcription patterns. Amongst other chromatin regulatory functions, many of these modifications are essential for cell differentiation and provide mechanisms for maintaining lineage-specific identity and cell functions through the life of an organism. Conversely, dysregulation of epigenetic modifications contributes to a wide range of diseases and syndromes, including congenital anomalies, cancer, diabetes and behavioural conditions (1-4).

The maternal and paternal genomes transmit genetic and epigenetic information to offspring at fertilisation. While oocyte and sperm chromatin are respectively organised in distinct histone and protamine-mediated structures, the vast majority of maternal and paternal alleles achieve epigenetic equivalence within a short period after fertilisation, a process that relies partly on proteins and RNAs that are maternally inherited in the oocyte. However, some genes maintain parent-specific epigenetic patterns that were established during sperm and oocyte development. In mice and humans, these genes include around 120 imprinted genes that are typically marked either by maternal or paternal DNA methylation, an epigenetic state that is transmitted to, and maintained in offspring and is essential for parent-of-origin specific gene regulation during development (5-8). While genomic imprinting provides an unequivocal example of epigenetic inheritance, evidence for other epigenetically inherited states that may affect biallelically expressed genes are rare and the mechanisms underlying such inheritance are poorly understood (5). Given the potential for epigenetic states to influence offspring development, identifying the specific chromatin-modifying complexes that epigenetically regulate developmental genes and may influence establishment of an appropriate epigenetic landscape in oocytes would enhance understanding of the mechanisms underlying inherited phenotypes and disease, and of how these mechanisms may contribute to evolution.

Histone 3 Lysine 27 trimethylation (H3K27me3) is a critical epigenetic modification catalysed by the Polycomb Repressive Complex 2 (PRC2). PRC2 contains three essential core protein subunits: Suppressor of Zeste 12 (SUZ12), Embryonic Ectoderm Development (EED) and Enhancer of Zeste 1/2 (EZH1/2), all of which are required for histone methyltransferase activity (9-12). While EZH2 can function in PRC2-independent roles, EED is only known to mediate methylation of H3K27 as an essential component of PRC2 (13-19). Specific examples include an essential role for EED in repressing a wide range of developmentally important genes in embryonic stem cells (ESCs) through its essential role in establishing H3K27me3 (12, 20). While EZH2 also plays a major role in the repression of the same genes, the closely related protein EZH1 acts in a partially redundant manner and contributes both to H3K27me3 enrichment and gene repression (12). In other contexts, EZH2 can directly methylate non-histone target proteins such as PLZF in B lymphocytes of the immune system, and GATA4 in mouse fetal cardiomyocytes *in vivo* (13, 17).

PRC2 also plays important roles in sperm and oocytes, and throughout development. *De novo* germline mutations in human *EED, EZH2* and *SUZ12* underlie Cohen-Gibson, Weaver and Imagawa-Matsumoto syndromes which are characterised by perinatal overgrowth, skeletal malformation and cognitive deficit (21-30). Multiple studies in mice indicate that EZH2 and EED act as maternal factor proteins and/or mRNA that are required in mature oocytes to regulate the establishment and maintenance of X-inactivation in pre-implantation embryos (31-34). In addition, PRC2 regulates DNA methylation-independent non-canonical imprinting in mouse oocytes, a process that involves H3K27me3-dependent programming and paternally-biased expression of up to 20 genes in pre-implantation embryos and five genes in extraembryonic ectoderm and placenta until embryonic day (E)9.5 (8, 35). Maternal deletion of *Eed* resulted in loss of H3K27me3 imprints, biallelic expression of H3K27me3-imprinted genes in pre-implantation embryos and extraembryonic ectoderm, transient ectopic X-inactivation and male-biased embryo loss (33, 34). Moreover, mouse offspring generated by somatic cell nuclear transfer (SCNT) are typically born large as a result of placental hyperplasia, a phenotype that is caused by loss of H3K27me3 imprinting primarily of *Slc38a4* and *Sfmbt2-*embedded micro-RNAs specifically in the placenta (36-38). Although H3K27me3 imprinting specifically affects the placenta, embryonic growth restriction was also observed in embryos derived from oocytes lacking EED, but the cause of this phenotype is not understood (33). While H3K27me3-dependent imprinting (non-canonical) has been recently identified, classical (or canonical) genomic imprinting is much more extensively studied and is generally considered to be mediated by DNA methylation (8, 39). Here, we refer to canonical DNA methylation-based genomic imprints as classical imprinting and non-canonical imprinting as H3K27me3-dependent imprinting.

We previously found that deletion of *Eed* in growing oocytes led to post-natal overgrowth of offspring, indicating that maternally-derived PRC2 mediates effects on offspring that were independent of maternal genetic inheritance (40). To understand the potential mechanisms underlying developmental outcomes in offspring from *Eed* null oocytes, we explored the role of PRC2 in oocytes. We demonstrate that EZH2, EED and SUZ12 are transiently expressed during the earliest stages of oocyte growth to establish H3K27me3 in the promoters of developmentally important genes in mice, and that H3K27me3 is conserved on many of these genes in human GV stage oocytes. In mice, PRC2 activity immediately preceded the upregulation of the essential *de novo* DNA methylation co-factor DNMT3L, indicating that patterning of PRC2 target genes precedes DNA methylation. While *Eed* repressed several imprinted genes in oocytes, 98% of the PRC2 target genes we identified were not imprinted, but were genes that regulate neurogenesis, haematopoiesis and other processes in tissue morphogenesis. These genes were not dysregulated in pre-implantation offspring and lost H3K27me3 during this period of development in wild type (*wt*) embryos.

## Results

### EZH2, EED and SUZ12 localise to chromatin during a discrete period of primary to secondary oocyte growth

Previous studies have provided varying reports of EED, EZH2 and SUZ12 in GV stage and mature oocytes, and zygotes (31, 41-43), but the stages at which all three core components of PRC2 are detected in growing oocytes have not been defined. To determine when PRC2 is detected in the nucleus or associated with chromosomes in growing, GV and MII oocytes, and in zygotes, we profiled EZH2, SUZ12 and EED throughout oocyte growth in *wt* mice using immunofluorescence (IF). EZH2 was detected in the oocyte nucleus of primordial to antral stage follicles, but SUZ12 and EED were detected only in primary and secondary follicle oocytes and not in primordial or antral stage oocytes (Fig. 1A). Notably, co-expression of EED, EZH2 and SUZ12 in primary-secondary follicle oocytes occurred immediately before the expression of DNMT3L (DNA methyltransferase 3-Like), which marks the onset of *de novo* DNA methylation in growing oocytes (Fig. 1B), consistent with the initiation of H3K27me3 in oocytes prior to DNA methylation (44). While EZH2 was detected in the nuclei of fully-grown surrounded nucleolus (SN) GV oocytes, SUZ12 and EED were not (Fig. 1C). Although PRC2 has been detected in the cytoplasm of mature metaphase II (MII) oocytes (41-43), EED, EZH2 and SUZ12 were not detected on the chromosomes of MII oocytes in this study (Fig. 1D). However, all three PRC2 components were readily detected in maternal and paternal pronuclei of zygotes approximately 12 hours (h) post-fertilization (Fig. 1E). As embryonic activity of PRC2 does not occur until the 4-cell to morula stage (31, 34), the rapid recruitment of PRC2 to the pronuclei may reflect a supply of cytoplasmic PRC2 proteins (41-43) or could be derived from mRNAs in the mature oocyte. Taken together, these data identify a transient window during which all three PRC2 components are present and may therefore contribute to PRC2-dependent epigenetic programming in primary-secondary oocytes, immediately before genome-wide *de novo* DNA methylation and prior to the formation of GV-oocytes.

**Fig 1.**
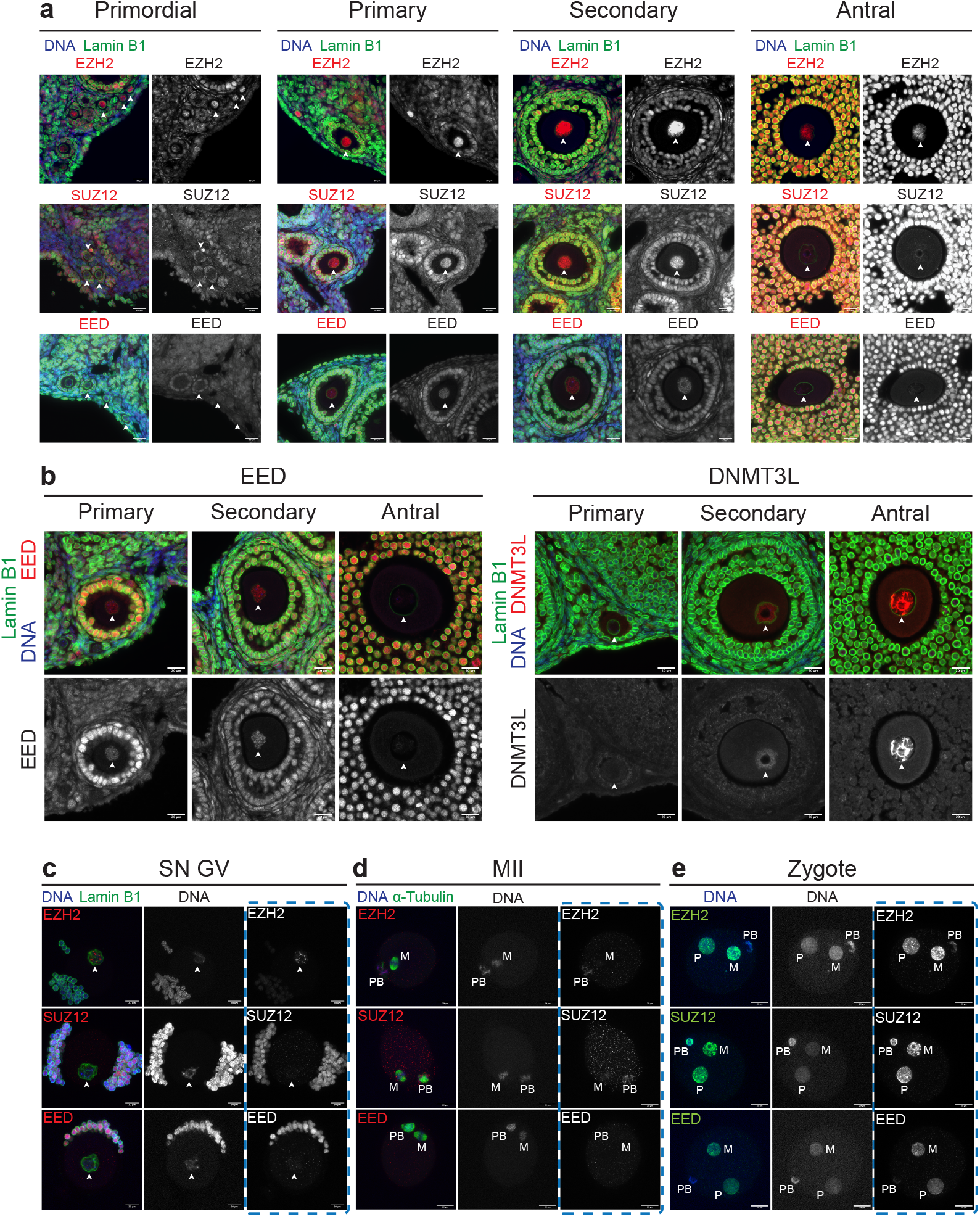
PRC2 acts transiently within primary and secondary follicle growing oocytes. **(a-d)** Representative images of EZH2, SUZ12 and EED (red) IF analysis in **(a)** primordial, primary, secondary and antral follicles. **(b)** Comparison of EED (red, left panels) versus DNMT3L (red, right panels) IF analysis in primary, secondary and antral follicles. **(c)** Surrounded nucleolus (SN) GV oocytes. **(d)** MII oocytes. α-Tubulin (green) identifies meiotic spindles. M: metaphase plate, PB: polar body. **(e)** Zygotes 12h after fertilisation for ≥ 10 zygotes imaged per antibody combination. M: maternal pronucleus, P: paternal pronucleus, PB: polar body. In **a-c** white arrowheads indicate the oocyte nucleus defined by Lamin B1 (green) and DAPI (blue) shows DNA. In **c-e** images represent compressed z-stack images of wholemount oocytes or zygotes. Scale bars: 20 μm. Images in **a** and **b** are representative of two ovaries from three separate females and in **c-e** images are representative of ≥10 oocytes per antibody combination.

### PRC2 is required for repression of developmental genes in growing oocytes

To investigate whether the transient activity of PRC2 in oocytes of primary-secondary follicles has functional importance, we deleted *Eed* using *Zp3Cre*, which leads to target gene excision specifically in oocytes from the primary follicle stage (40, 45). Mating *Eed*^*fl/fl*^ females and *Eed*^*fl/+*^;*Zp3Cre* males yielded *Eed*^*fl/fl*^ (*Eed-wt*), *Eed*^*fl/+*^;*Zp3Cre* (*Eed-het*) and *Eed*^*fl/fl*^;*Zp3Cre* (*Eed-hom*) females. Following deletion of *Eed*, H3K27me3 was reduced by 35% in primary follicles of *Eed-hom* females (Fig. 2A-B). Depletion of H3K27me3 continued in secondary follicles (Fig. 2A, C), and was almost completely lost in fully-grown GV oocytes, with 85% and 93% reductions in global H3K27me3 in *Eed-hom* oocytes compared to *Eed-wt* at these stages respectively (Fig. 2A, 2C, 3A-B).

**Fig 2.**
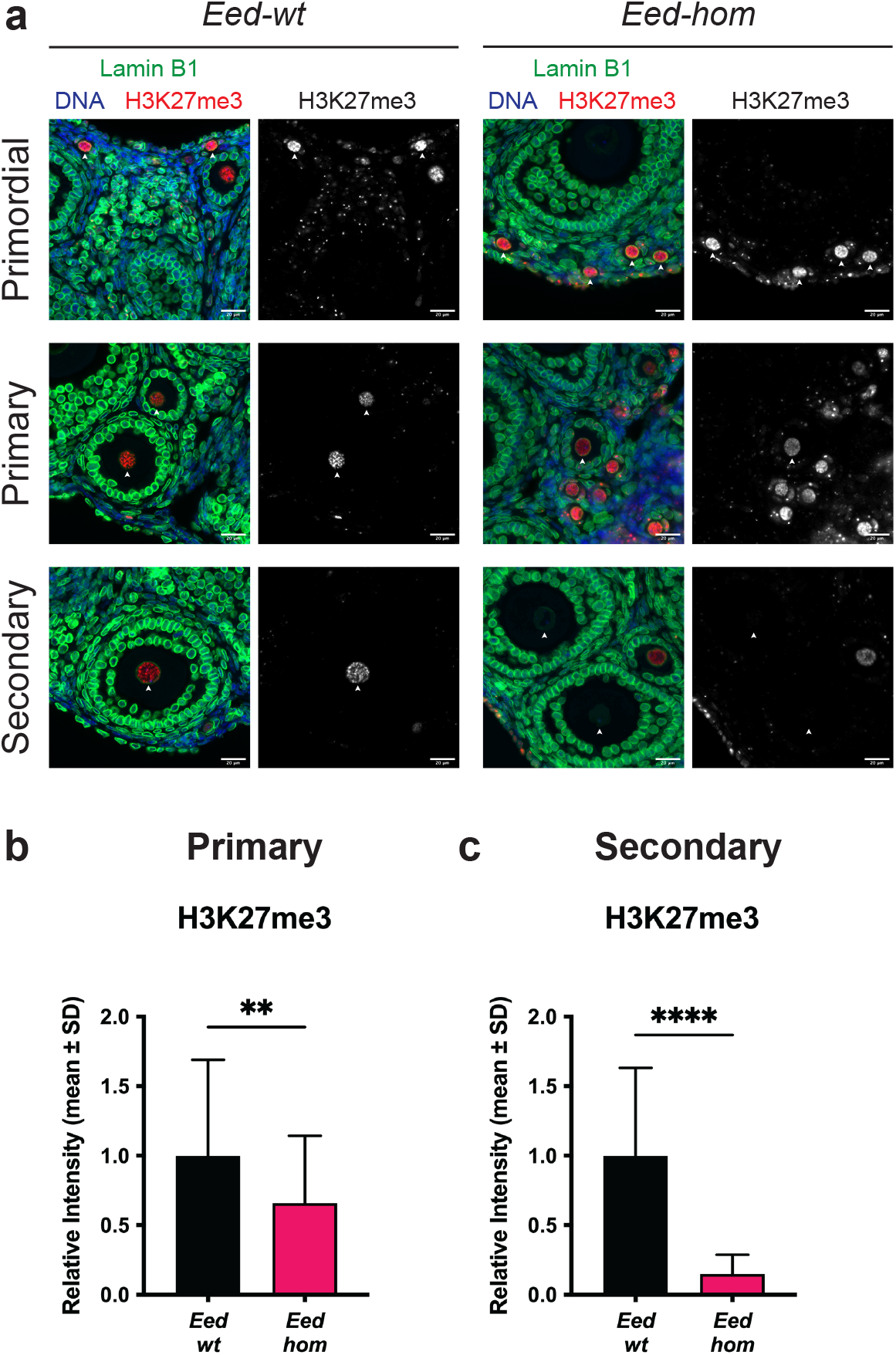
Deletion of *Eed* in oocytes reduced H3K27me3 in oocytes of primary and secondary follicles. **(a)** Representative images of H3K27me3 (red) immunostaining analysis in primordial (top), primary (middle) and secondary (bottom) follicle oocytes from *Eed-wt* and *Eed-hom* females. White arrowheads indicate the oocyte nucleus as defined by Lamin B1 (green). DAPI (blue) shows DNA in somatic cells. Images are representative of two ovaries from three biological replicates. Scale bars: 20 μm. **(b-c)** Quantification of H3K27me3 within oocyte nuclei of primary **(b)** and secondary **(c)** follicles from *Eed-wt* and *Eed-hom* females. Average intensity of *Eed-wt* was set to 1.0. \*\**P* < 0.005, two-tailed Mann-Whitney U test, N = 63 *Eed-wt* and 67 *Eed-hom* primary follicle oocytes. \*\*\*\**P* < 0.0001, two-tailed Mann-Whitney U test, N = 45 *Eed-wt* and 47 *Eed-hom* secondary follicle oocytes. Error bars represent mean ± standard deviation.

**Fig 3.**
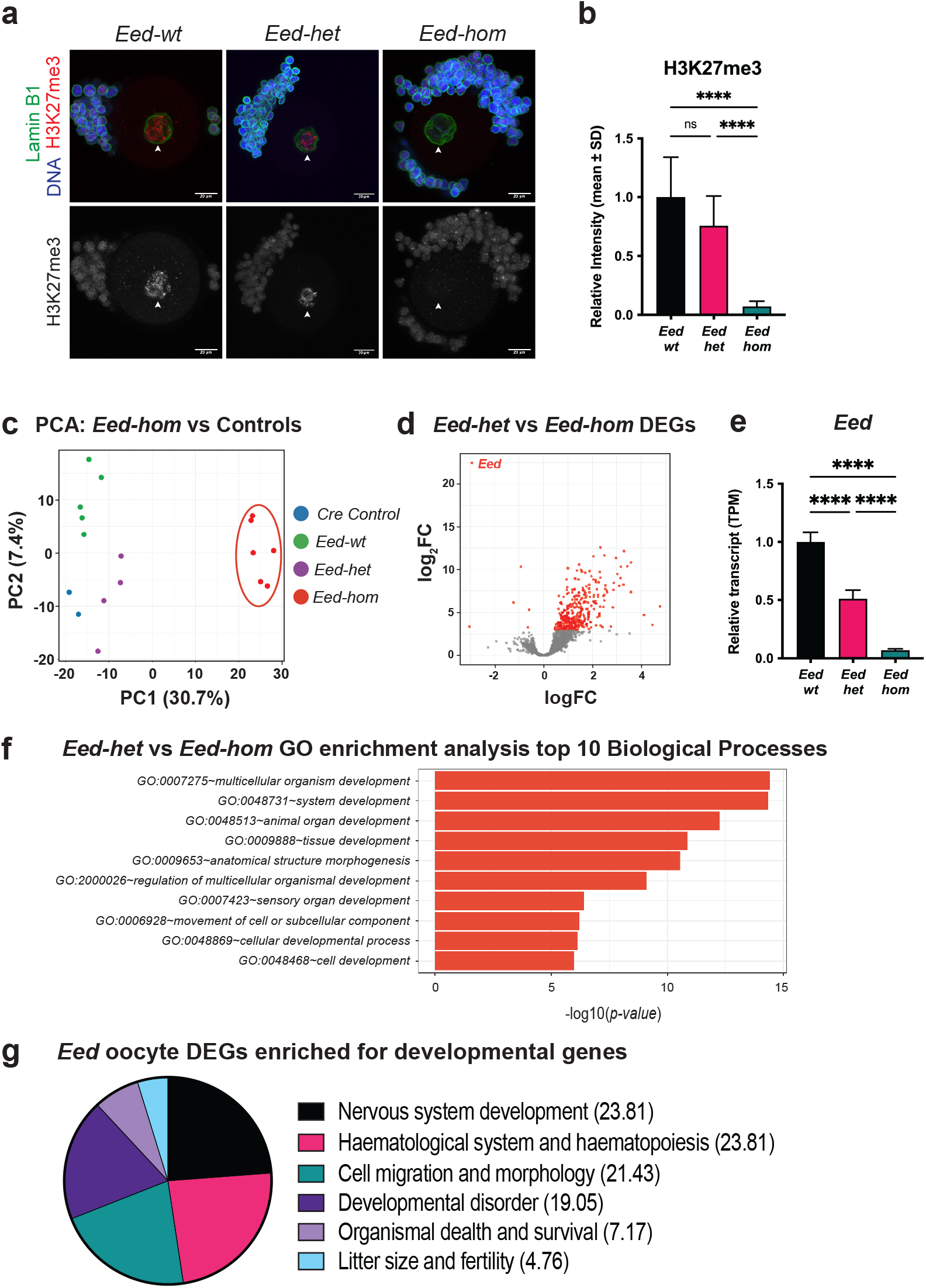
*Eed* is required for H3K27me3 establishment and developmental gene silencing in growing oocytes. **(a,b)** Representative images **(a)** and quantification **(b)** of H3K27me3 (red) IF in *Eed-wt, Eed-het*, and *Eed-hom* SN GV oocytes. White arrowheads indicate the oocyte nucleus as defined by Lamin B1 (green). DAPI (blue) shows DNA in somatic cells. Images represent 3-4 females per genotype, with 16-21 oocytes imaged per genotype. Scale bars: 20 μm. Average intensity of *Eed-wt* was set to 1.0. \*\*\*\**P* < 0.0001, Kruskall-Wallis test plus Dunn’s multiple comparisons test, error bars represent mean ± standard deviation. **(c)** Principal Component Analysis (PCA) of RNAseq data for *Eed-hom* (n=6) vs *Eed-het* (n=4), *Eed-wt* (n=5) and *Eed-wt Cre* (n=2) controls. **(d)** Differential gene expression analysis of *Eed-het* vs *Eed-hom* oocytes represented by volcano plot showing logFC against statistical significance. Genes with FDR-adjusted *P* < 0.05 are coloured in red. Deletion of *Eed* resulted in 349 significant DEGs (*Eed* oocyte DEGs), with 343 genes upregulated and 6 genes downregulated, including *Eed*. **(e)** Relative *Eed* transcript levels (transcripts per million reads; TPM) in *Eed-wt, Eed-het* and *Eed-hom* GV oocytes. Average expression of *Eed-wt* was set to 1.0. **(f)** GO enrichment analysis of *Eed* oocyte DEGs representing the top 10 significantly different biological processes impacted. **(f)** Pie chart displaying the proportion of significant pathways identified using Ingenuity pathway analysis

To determine how loss of H3K27me3 impacted oocyte transcription, we collected *Eed-wt, Eed-het, Eed-hom* and *Eed*^*+/+*^;*Zp3Cre* (*Eed*-*wt Cre* control) fully-grown SN GV oocytes and performed RNA-seq. The proportion of SN GV oocytes in *Eed-hom* females was 65% compared to 58% in *Eed-wt* females (Supp. Fig. 1), demonstrating that loss of EED and H3K27me3 did not detrimentally affect formation of fully-grown oocytes. Principal component analysis revealed that *Eed-hom* oocytes were transcriptionally distinct from *Eed-het* and *Eed*-*wt* oocytes (Fig. 3C). Further analysis identified 349 genes that we termed *Eed* oocyte Differentially-Expressed Genes (DEGs) as they were differentially expressed between *Eed-hom* and *Eed-het* oocytes (FDR<0.05; Fig. 3D; Supp. Table 1). Strikingly, 98% (343 genes) of the *Eed* oocyte DEGs were derepressed, and only 2% (six genes), including *Eed*, were downregulated (Fig. 3D-E; Supp. Table 1). H3K27me3 levels were not different and only two *Eed* oocyte DEGs (*Mt1* and *Exoc)* as well as *Eed* were identified between *Eed-wt* and *Eed-het* oocytes (Fig. 3A-B), indicating that EED function in *Eed-wt* and *Eed-het* oocytes was similar. As EED protein was only detected in primary-secondary oocytes prior to the GV stage, these data strongly indicate that PRC2 establishes a repressive state in primary and secondary follicle oocytes that is maintained in GV oocytes.

Gene ontology (GO) and ingenuity pathway analyses (IPA) revealed that the *Eed* oocyte DEGs were strongly associated with fetal development, including the IPA categories of nervous system development (23.81%), haematopoiesis (23.81%), cell migration and morphology (21.43%) and developmental disorders (19.05%; Fig. 3F-G). Several genes involved in bone development, including *Prrx, Gli2, Sox5, Sox6, Hoxd9, Hoxd13, Bmp7, Sik3* and *Dcn* were also de-repressed (Supp. Table 1). The neurogenesis and bone developmental genes are of interest as impaired skeletal and cognitive development are prominent features of Cohen-Gibson syndrome which results from *de novo* germline mutations in *EED* (21-23, 26). Although 4.67% of the *Eed* oocyte DEGs were associated with “litter size and fertility”, categories associated with oocyte or ovarian development were not represented (Fig. 3G). Moreover, similar numbers of oocytes in *Eed-hom, Eed-het* and *Eed-wt* females indicated that oocyte growth and formation of fully-grown oocytes was not impeded (Supp. Fig. 1). Together, these observations strongly suggested that PRC2 establishes repressive H3K27me3 on a wide range of developmentally important genes during oocyte growth, >95% of which were not primarily involved in oogenesis. Surprisingly, comparison of the *Eed* oocyte DEGs with 209 transcription factors identified as direct target genes of PRC1 and PRC2 in embryonic stem cells (20) identified only 8 common genes (*Hoxd9, Hoxd13, Otx1, Lhx2, Six1, Nr2f2, Ovol1* and *Nfatc1*), indicating that the genes that were de-repressed on *Eed* null oocytes were not typical Polycomb target genes in ESCs (Supp Table 2).

### PRC2 regulates establishment of H3K27me3 on developmental genes in growing oocytes

To determine whether H3K27me3 was normally present in the promoters of *Eed* oocyte DEGs in GV oocytes we compared the *Eed* oocyte DEGs to H3K27me3 Chromatin Immunoprecipitation-sequencing (ChIP-seq) datasets from wild type mouse GV and MII oocytes from Zheng and colleagues and Liu and colleagues (44, 46). Of 349 *Eed* oocyte DEGs, the majority were identified both in the Zheng and Liu datasets (328 in Zheng, and 312 in Liu). Of the 328 genes from Zheng *et al*., 111 (34%) and 127 (39%) had H3K27me3 peaks in GV and MII oocyte datasets, and 169 (52%) had H3K27me3 peaks in the Liu MII dataset (Supp Table 3). Comparison of all datasets (our *Eed* oocyte DEGs with the Zheng GV and MII and Liu MII datasets) identified 99 DEGs with H3K27me3 in all three ChIP-seq datasets, which we defined as ‘high confidence H3K27me3-enriched oocyte DEGs (Fig. 4A; Supp. Table 4). Of these 99 genes, comparison with data from a study of H3K27me3 in sperm (47) revealed that 88 also carried H3K27me3 on the paternal allele indicating that the majority of these genes are subject to H3K27 methylation in both the male and female germlines (Supp. Table 4). While only ∼30% of the *Eed* oocyte DEGs contained H3K27me3 under these highly stringent criteria, we propose that this is a conservative estimate given that low input ChIP-seq data were used, potentially limiting sensitivity compared to RNA-seq analysis. Supporting this, 114 H3K27me3-enriched DEGs overlapped between the Liu and Zheng MII datasets, indicating that not all genes with H3K27me3 were consistently detected in the ChIP-seq analyses.

**Fig 4.**
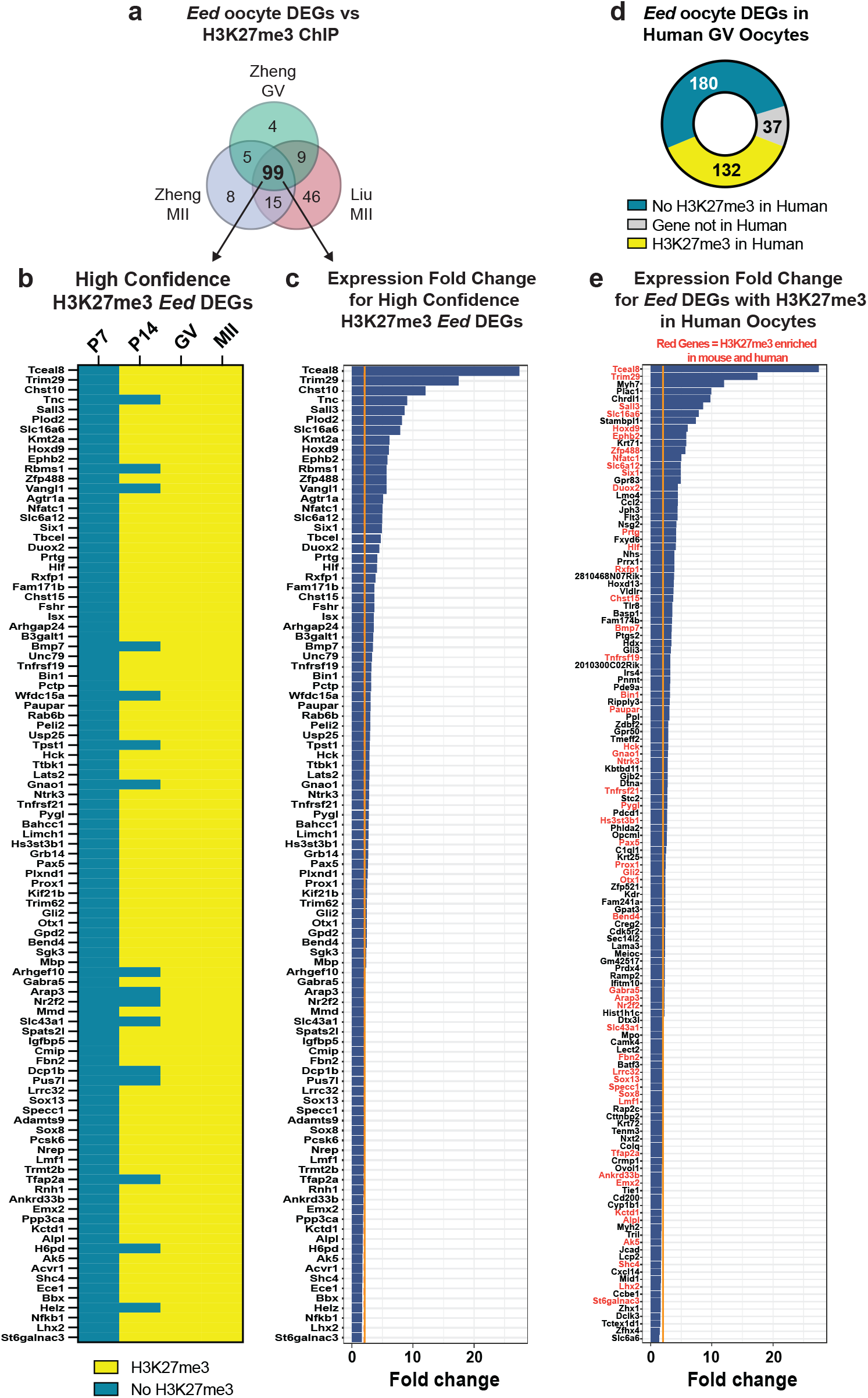
H3K27me3 is established on *Eed* oocyte DEGs in primary – secondary mouse oocytes and is conserved in human GV oocytes. **(a)** Venn diagram showing *Eed* oocyte DEGs that contained H3K27me3 promoter peaks in GV and MII oocyte H3K27me3 ChIP-seq datasets (44, 46) identifying 99 ‘high confidence’ H3K27me3 enriched *Eed* oocyte DEGs. **(b)** Heat map showing promoter H3K27me3 enrichment status of 99 high confidence H3K27me3 enriched *Eed* oocyte DEGs identified in P7, P14, GV and MII oocyte H3K27me3 ChIP-seq datasets from Liu *et al*., and Zheng *et al*., (44, 46). Blue: No H3K27me3 peaks, yellow: indicates presence of H3K27me3 peaks. **(c)** Expression fold change of the 99 high confidence H3K27me3 enriched *Eed* oocyte DEGs in *Eed-hom* oocytes relative to *Eed-het*. Orange line indicates two-fold change **(d)** Donut chart showing the promoter H3K27me3 enrichment status of *Eed* oocyte DEGs in Human GV oocytes (48). Grey: Not conserved in humans, blue: no H3K27me3 peaks in promoter, yellow: H3K27me3 peak present in promoter. **(e)** Expression fold change of 132 mouse *Eed* oocyte DEGs that were H3K27me3 enriched in human GV oocytes. *Eed* oocyte DEGs commonly enriched for H3K27me3 in human and mouse GV oocytes are marked with genes names in red. Orange line indicates two-fold change. For **(a,b and d)**, promoter region was defined as 2000bp upstream and downstream of TSS, overlap of > 200bp H3K27me3 peaks with the promoter region was considered H3K27me3 enriched.

Gradual enrichment of H3K27me3 has been demonstrated in growing oocytes (44). Using these additional ChIP-seq datasets (44), we next investigated at what stage H3K27me3 peaks were established at the high confidence H3K27me3-enriched oocyte DEGs by determining the H3K27me3 state at the promoter regions of these genes in post-natal day (P)7 (primary) and P14 (secondary) growing oocytes. None of the 99 high confidence H3K27me3-enriched oocyte DEGs had H3K27me3 peaks in primary (P7) oocytes, but 83 had H3K27me3 peaks in secondary (P14) oocytes and all 99 DEGs had H3K27me3 in GV and MII oocytes (Fig. 4B; Supp. Table 4). Accordingly, each of these genes were upregulated in *Eed-hom* GV oocytes (Fig. 4C). Collectively, these data demonstrate that H3K27me3 was established within the promoters of developmental genes during the window in which all three core components of PRC2 were detected in primary-secondary oocytes and that EED was required for their repression given that these genes were de-repressed in *Eed*-null GV oocytes.

### PRC2 establishment of H3K27me3 at developmental genes is conserved in human oocytes

To understand whether the mouse *Eed* oocyte DEGs were also enriched for H3K27me3 in human oocytes, we examined published H3K27me3 data from human GV oocytes (48), defining promoter regions 2000bp upstream – 2000bp downstream of the TSS as we did for the mouse datasets. Of 349 *Eed* oocyte DEGs, 37 were excluded as ‘not conserved in human’, including 25 predicted genes, RIKEN transcripts or pseudogenes (Fig. 4D; Supp. Table 3). Of the 312 remaining *Eed* oocyte DEGs, 132 contained H3K27me3 in their promoters in human GV oocytes (Fig 4D; Supp. Table 3; Supp. Table 5). Of the 132 *Eed* oocyte DEGs containing H3K27me3 in human GV oocytes, 79 and 54 also contained H3K27me3 in the mouse MII and GV datasets generated by Liu *et al*. and Zheng *et al*., respectively (Supp. Table 5 (44, 46)). Moreover, all 132 *Eed* oocyte DEGs identified as H3K27me3 enriched in human were upregulated in *Eed-hom* oocytes (Fig. 4E). In common with the lack of nuclear-localised EED or SUZ12 protein in mouse GV oocytes (Fig. 1C), human GV oocytes lack *EED* and *SUZ12* transcripts (48). Together, with the observation that *Eed* oocyte DEGS are almost exclusively de-repressed with the loss of EED and H3K27me3, these data strongly indicate that H3K27me3 establishment occurs on *Eed* oocyte DEGs prior to the GV stage in both human and mouse growing oocytes.

### *Eed* oocyte DEGs include non-imprinted autosomal, imprinted and X-linked genes

As EED regulates DNA methylation-independent H3K27me3 imprinting in oocytes, we compared the *Eed* oocyte DEGs against a previously published list of 76 putative H3K27me3-imprinted genes (35). Of the 349 *Eed* oocyte DEGs we identified in GV oocytes, five (*Bbx, Bmp7, Rbms1, Sall3* and *Prox1*) were putative H3K27me3 imprinted genes (Fig. 5A; Supp. Table 6). These genes were all upregulated in *Eed-hom* oocytes (Fig. 5B), demonstrating that EED is required for their repression in oocytes. Of interest, analysis of a sperm ChIP-seq dataset (47) revealed that all five genes also contained H3K27me3 on the paternal allele, indicating that these genes have similar H3K27me3 signatures in male and female gametes (Supp. Table 3). Of these five genes, paternally biased expression was observed for three in androgenetic morula (*Bbx, Bmp7* and *Rbms1*) and one in blastocysts (*Rbms1*) (35). Notably, Inoue *et al*. identified five H3K27me3-imprinted genes (*Gab1, Phf17, Sfmbt2, Slc38a4 and Smoc1*) that maintain paternal biased expression in epiblast, visceral endoderm, extra-embryonic ectoderm and/or E9.5 placenta (35). Loss of imprinting at either *Slc38a4* or micro-RNAs within *Sfmbt2* has been functionally associated with placental hyperplasia in SCNT derived offspring (36, 38). In addition, *Smoc1* and *Gab1* have also been implicated in this phenotype (37). However, none of these genes were *Eed* oocyte DEGs (Supp. Table 6).

**Fig 5.**
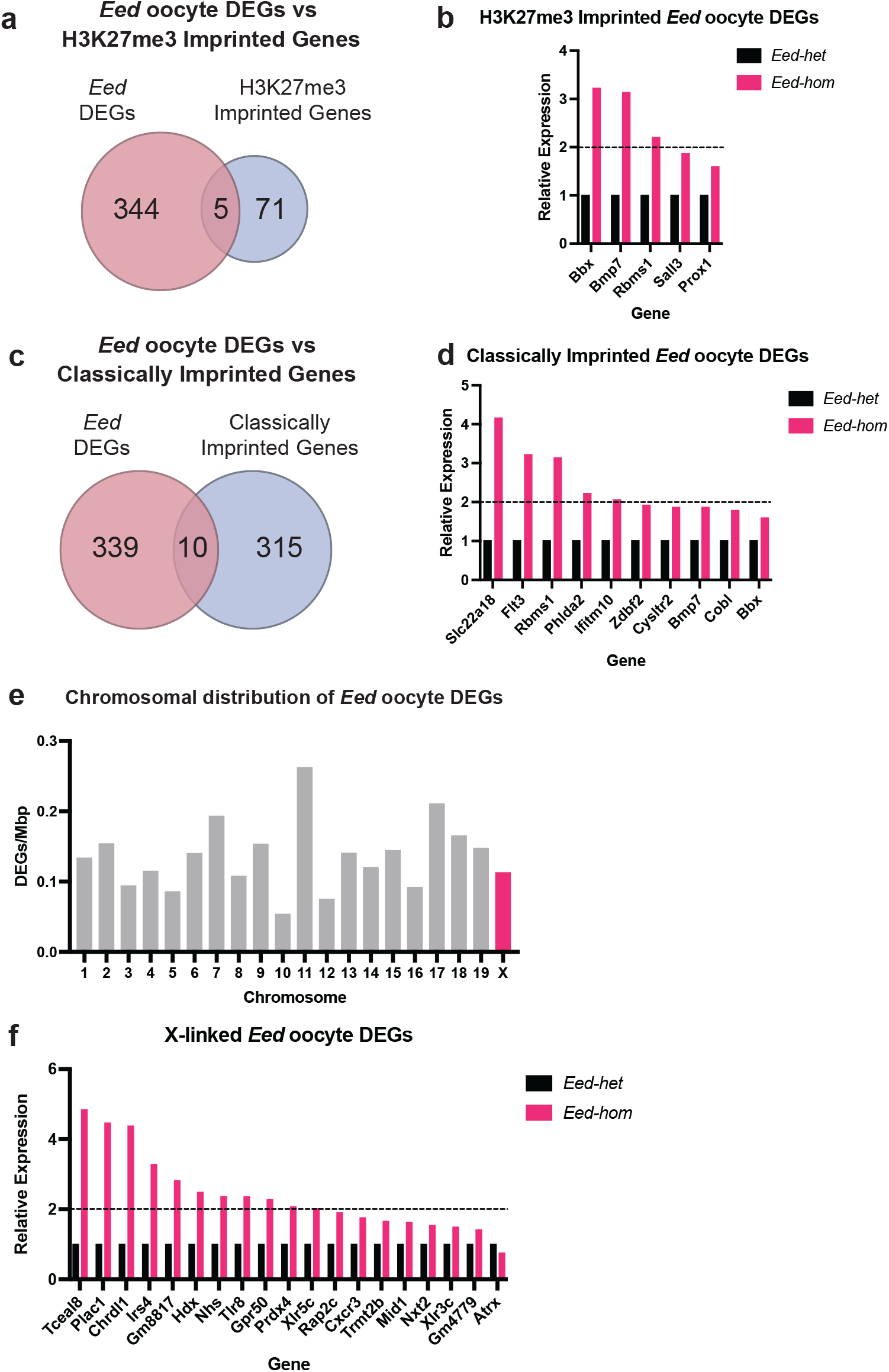
EED is required for repressing a wide range of genes in growing oocytes that are not canonically or non-canonically imprinted or X-linked genes. **(a)** Venn diagram comparing *Eed* oocyte DEGs against putative H3K27me3 imprinted genes (35). **(b)** Expression of putative H3K27me3 imprinted *Eed* oocyte DEGs in *Eed-hom* oocyte relative to *Eed-het*. Data represent the mean transcripts per million (TPM), with *Eed-het* mean set to 1.0. **(c)** Venn diagram comparing *Eed* oocyte DEGs against known or predicted classically imprinted genes (49, 50). **(d)** Expression of known or predicted classically imprinted *Eed* oocyte DEGs in *Eed-hom* oocyte relative to *Eed-het*. Data represent the mean transcripts per million (TPM), with *Eed-het* mean set to 1.0. **(e)** graphical representation of *Eed* oocyte DEGs per mega base vs chromosome for autosome and the X-chromosome. **(f)** Expression of X-linked *Eed* oocyte DEGs in *Eed-hom* oocyte relative to *Eed-het*. Data represent the mean transcripts per million (TPM), with *Eed-het* mean set to 1.0.

In order to determine whether any *Eed* oocyte DEGs were imprinted genes, we also compared the 349 *Eed* oocyte DEGs to 325 known or predicted classically imprinted genes in mice (49, 50). Ten *Eed* oocyte DEGs were identified as imprinted genes (Fig. 5C; Supp. Table 6; *Zdbf2, Ifitm10, Phlda2, Rbms1, Bmp7, Bbx, Flt3, Slc22a18, Cobl* and *Cysltr2*). As with the putative H3K27me3-imprinted genes, these were all upregulated in *Eed-hom* oocytes (Fig. 5D) indicating that EED is required to silence these genes in oocytes. Analysis of the sperm dataset (47) revealed that all of these genes contained H3K27me3 on the paternal allele in sperm and three of these genes (*Bbx, Bmp7* and *Rbms1*) were also included on the H3K27me3-imprinted gene list (Supp. Table 3). In human GV oocytes *Bmp7, Sall3, Prox1, Zdbf2, Phlda2, Flt3* and *Ifitm10* also carried H3K27me3, but *Bbx, Rbms1, Cysltr2* and *Slc22a4* did not.

In mice, X-inactivation is initiated after fertilisation from the 2/4 cell stage and is restricted to the paternal X-chromosome in pre-implantation offspring prior to establishment of random X-inactivation in embryonic cells after implantation. While PRC2 regulates X-inactivation in pre-implantation embryos and somatic cells of post-implantation embryos, the inactive X is reactivated in XX primordial germ cells and both X-chromosomes are active in growing oocytes (51-53). To determine whether there was any bias in gene silencing across the autosomes and the X-chromosome in EED-deficient oocytes we determined the relative representation of the *Eed* oocyte DEGs across all chromosomes. However, representation of the oocyte DEGs across the autosomes and X-chromosome was similar, with 19 of 349 genes X-located and no substantial bias towards the X-chromosome or particular autosomes (Fig. 5E; Supp. Table 7). As with most genes on the autosomes, of the 19 X-linked genes identified, 18 were upregulated in *Eed-hom* oocytes (Fig. 5F) demonstrating that EED and H3K27me3 contribute to silencing individual X-linked genes in the absence of X inactivation in oocytes.

Together, these data indicated that of the 349 *Eed* oocyte DEGs identified, 12 were imprinted genes, 19 were located on the X-chromosome and the remaining 328 were non-imprinted autosomal genes, many of which are known to regulate development.

### LINE-1 transposable elements were not de-repressed in *Eed*-null oocytes

Previous studies have proposed and/or demonstrated a link between H3K27me3 and the repression of transposable elements (TEs) in fetal germ cells and embryonic stem cells when DNA methylation levels are low during epigenetic reprogramming (54-57). To determine whether transcription of transposable elements was affected in oocytes lacking EED, we demasked repeat sequences in our RNA-seq data and analysed the expression of LINE-1 (L1) elements. The total input reads indicated that similar percentages of L1 element reads aligned in *Eed-hom, Eed-het* and *Eed-wt* oocytes (Supp. Fig. 2A). Moreover, regardless of whether elements mapped uniquely or multi-mapped, read totals indicated similar expression levels for L1s in oocytes of all genotypes (Supp. Fig. 2B). Analysis of the extent to which multiple reads occurred revealed that the majority of reads mapped 1, 2 or 3 times (accounting for ∼80% of all reads), while around 10% mapped 4-5 times and the remaining reads mapped 5-20 times, with no differences in mapping between genotypes (Supp. Fig. 2C). Principal component analysis revealed overlapping clusters for samples for all genotypes and no differentially expressed L1 elements were identified based on a threshold of FDR<0.05 (Fig. 6A, B). Together these data indicate that loss of H3K27me3 in GV oocytes did not substantially alter L1 expression.

**Fig 6.**
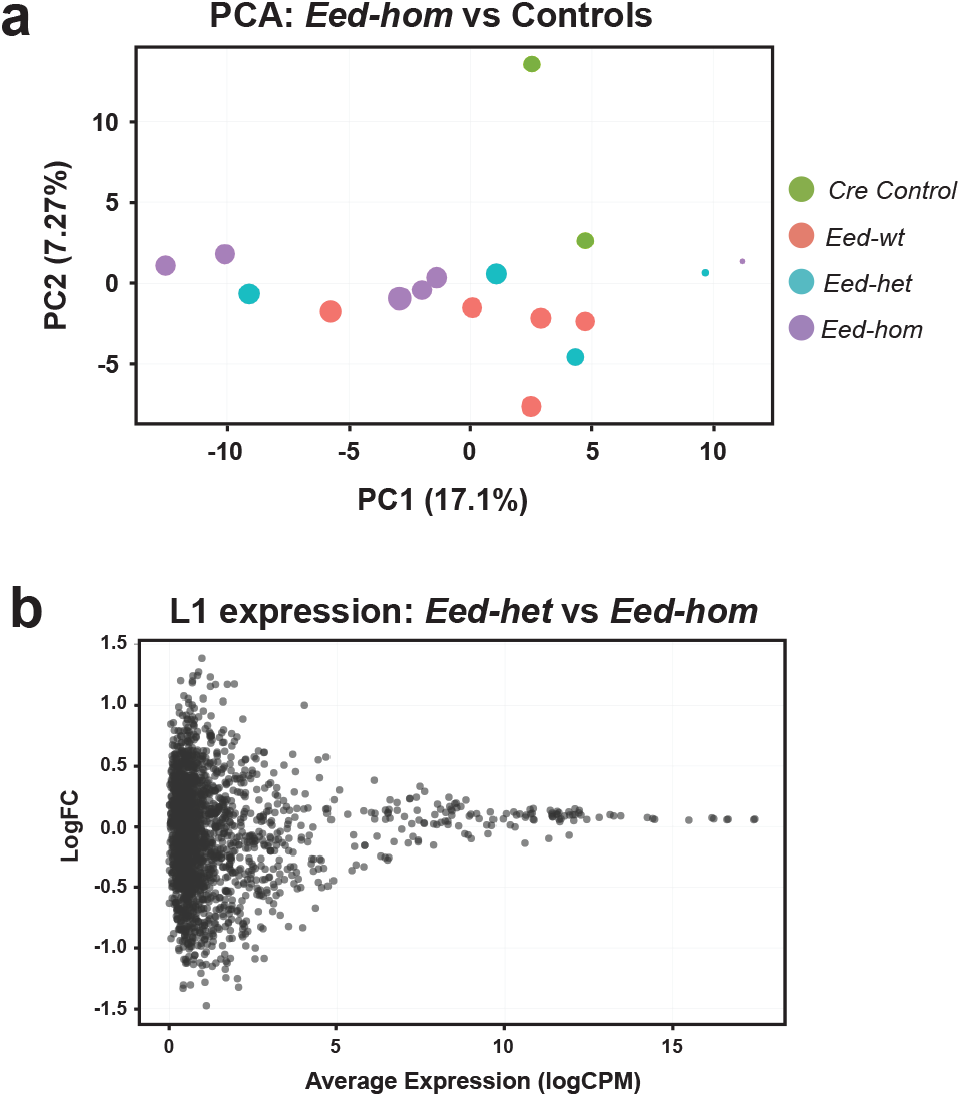
Loss of *Eed* in growing oocytes did not impact expression of LINE-1 transposons. **(a)** Principal Component Analysis (PCA) of RNA-seq data for L1 elements in *Eed-hom* (n=6) vs *Eed-het* (n=4), *Eed-wt* (n=5) and *Eed-wt Cre* (n=2) controls. **(b)** Differential expression analysis of L1 elements for *Eed-het* versus *Eed-hom* females.

### *Eed* oocyte DEGs were not dysregulated in pre-implantation embryos

Our observations revealed a highly specific window during which all three components of PRC2 were present in primary-secondary oocytes and identified a role for EED in establishing H3K27me3 on a wide range of developmentally important genes in primary-secondary stage oocytes. As >95% of these genes are not known to regulate oocyte development, yet EED is required for their repression in oocytes, we proposed that loss of EED-dependent repression of these genes in oocytes may result in their de-repression in pre-implantation embryos. To determine if this was the case we re-analysed RNA-seq data from morula and blastocysts derived from *Gdf9Cre Eed* null (33) and *Zp3Cre Eed* null oocytes (34). The *Eed-*deleted oocyte-derived morulae contained 128 DEGs (P<0.05, all with FDR∼1.0; Supp. Table 8) and the blastocysts contained 400 DEGs (P<0.05; FDR<0.05; Supp. Table 9) compared to their *Eed*-*wt* oocyte-derived counterparts. Six *Eed* oocyte DEGs were dysregulated in morulae (*Plxnd1, Tceal8, Rap2c, Bbx, Xlr3c* and *Trmt2b;* Supp. Fig. 3), and five were dysregulated in blastocysts (*Chrdl1, Lonrf2, Trim6, Cyp1b1* and *Ccbe1*; Supp. Fig. 3). With the exception of *Tceal8* and *Bbx*, all were downregulated in pre-implantation embryos derived from *Eed-*null oocytes (Supp. Tables 8-9). Only two genes were commonly downregulated in the morulae and blastocyst datasets (*Tspan6* and *Gk*) and no genes were dysregulated in all three datasets (Supp. Fig. 3). Similarly, there were no genes in the morula DEGs and only seven genes (*Zfpm2, Pax9, Tbx4, Foxc2, Atoh8, Msx1* and *Vsx1*) in the blastocyst DEGs that were common with 209 genes identified as direct Polycomb target transcription factors in ESCs (Supp. Table 10; (20)).

To determine whether H3K27me3 was maintained at *Eed* oocyte DEGs in pre-implantation embryos, we used CUT&RUN data revealing the H3K27me3 state on the maternal and paternal alleles in morula stage embryos (Supp. Table 3) (33). Six *Eed* oocyte DEGs contained H3K27me3 on the maternal, but not the paternal allele in morula. Of the remaining 343 *Eed* oocyte DEGs, 322 were devoid of H3K27me3 on both the maternal and paternal alleles. The maternal and paternal alleles were not distinguishable for 21 genes in the source dataset (33). These data indicate that H3K27me3 is normally cleared from the vast majority of *Eed* oocyte DEGs in morula stage embryos.

## Discussion

In this study, we identified a discrete period during which all three core components of PRC2 co-localised to the nucleus in primary-secondary oocytes, that H3K27me3 was established on a wide range of developmental genes in this window and that EED was required to repress these genes in GV oocytes. This transient activity of PRC2 facilitated H3K27me3 establishment on a wide range of *Eed* oocyte DEGs immediately prior to DNMT3L upregulation, indicating that EED-dependent programming precedes *de novo* DNA methylation and that epigenetic programming is highly temporally and spatially regulated during oocyte growth. As many of the same genes were marked by EED-dependent H3K27me3 in mouse and human GV oocytes, the role of PRC2 appears to be conserved in human and mouse oocytes. These findings broaden the understanding of the temporal, spatial and functional activity of PRC2 in the female germline

The *Eed* oocyte DEGs identified included five H3K27me3 imprinted and seven classically imprinted genes (three of the classically imprinted genes detected were also listed as H3K27me3 imprinted genes). However, the vast majority of *Eed* oocyte DEGs were not known imprinted loci, but included many genes known to regulate cell differentiation during fetal development. Despite this, the *Eed* oocyte DEGs identified were not over-expressed in pre-implantation offspring and lost H3K27me3 during pre-implantation development. While the significance of PRC2 regulation of these genes requires further investigation, previous studies revealed that loss of EED in the oocyte affects early development and post-natal outcomes in opposing ways: offspring from *Eed* null oocytes exhibit both placental and offspring growth restriction at E10.5 (33), but were subsequently overgrown by early post-natal stages (40). While the mechanisms remain unclear, the collective data indicate that PRC2 acts at multiple levels of oocyte growth and pre-implantation development to modulate outcomes in offspring.

In previous studies there has been a significant focus on PRC2 as a maternal factor complex that regulates aspects of pre-implantation development, including X-inactivation (31-34, 58). In this study we detected EZH2 in oocytes from the primordial follicle stage through to the SN GV oocyte stage, whereas SUZ12 and EED were only identified within primary and secondary follicle stage oocytes. Since all three components are required for PRC2 methyltransferase activity (9-12), this identifies a discrete window in primary-secondary follicle oocytes during which PRC2 has the capacity to catalyse methylation of H3K27. While proteins for EED or SUZ12 were not detected in association with chromatin of GV and MII oocytes in our study, all three proteins were robustly detected in both maternal and paternal pronuclei 12 hours post-fertilisation, highlighting a role for maternally inherited PRC2 in zygotes. As previous reports detected SUZ12, EED and EZH2 protein in MII oocytes using western blots (41, 42), it is likely that a maternal supply of protein resides in the cytoplasm but is difficult to detect using IF. However, it is also possible that maternal RNA for *Eed, Suz12* and *Ezh2* is inherited via the oocyte and is translated to facilitate enrichment of PRC2 in the maternal and paternal pronuclei of zygotes. Regardless of the mode of inheritance, these observations suggest that PRC2 has distinct activities that differentially impact epigenetic regulation in growing oocytes and affect pre-implantation development in offspring.

Consistent with this, loss of EED in the oocyte affected different genes sets in oocytes and in pre-implantation offspring. While <5% of *Eed* oocyte DEGs were imprinted genes, the vast majority were not imprinted and many were associated with post-implantation cell differentiation and tissue development. Moreover, although EED was required for repression of these genes in oocytes, very few *Eed* oocyte DEGs were dysregulated in morula and blastocysts derived from *Eed-*null oocytes, and H3K27me3 was lost on these genes during pre-implantation development. While this is consistent with previous findings that many developmental genes lose H3K27me3 during pre-implantation development (44), it also indicates that their repression at this stage does not require H3K27me3. Despite this, PRC2 is required in the oocyte for regulating a large cohort of genes in pre-implantation embryos that are distinct from the *Eed* DEGs identified here, and are dysregulated when *Eed* is deleted in oocytes. Thus, while EED is required for enriching H3K27me3 and repressing a large cohort of developmental genes in primary-secondary oocytes, it also regulates a distinct set of genes in pre-implantation embryos. Therefore, PRC2 functions both to epigenetically program a wide range of imprinted and non-imprinted genes in oocytes and acts as a maternal factor in the zygote.

Our data indicates that H3K27me3 was established in the promoters of *Eed* oocyte DEGs immediately before the onset of DNA methylation, raising the possibility that H3K27me3 could influence the establishment of other epigenetic modifications in oocytes. While H3K27me3 and DNA methylation are generally mutually exclusive (44), H3K36me3 coincides with DNA methylation in oocytes (59) and H3K36me3 deposition increases in regions that subsequently acquire DNA methylation in oocytes (60). Further, while loss of H3K36me3 in oocytes results in ectopic H3K27me3 deposition in oocytes (59), it is not known whether H3K36me3 is altered in oocytes following loss of H3K27me3. As the relationship between DNA methylation/H3K36me3 and H3K27me3 is antagonistic, a potential role of H3K27me3 in oocytes may be to act as a “place keeper” that ensures the promoters of EED-dependent oocyte genes are not subject to other forms of epigenetic alteration (Fig. 7), such as DNA methylation or H3K36 methylation (59). Such an effect may be similar to that proposed for H3K27me3 in protecting regions from DNA methylation during sperm maturation (47) and may explain both the enrichment of H3K27me3 on genes that regulate fetal development and why H3K27me3 is cleared from these genes during pre-implantation offspring development.

**Fig 7:**
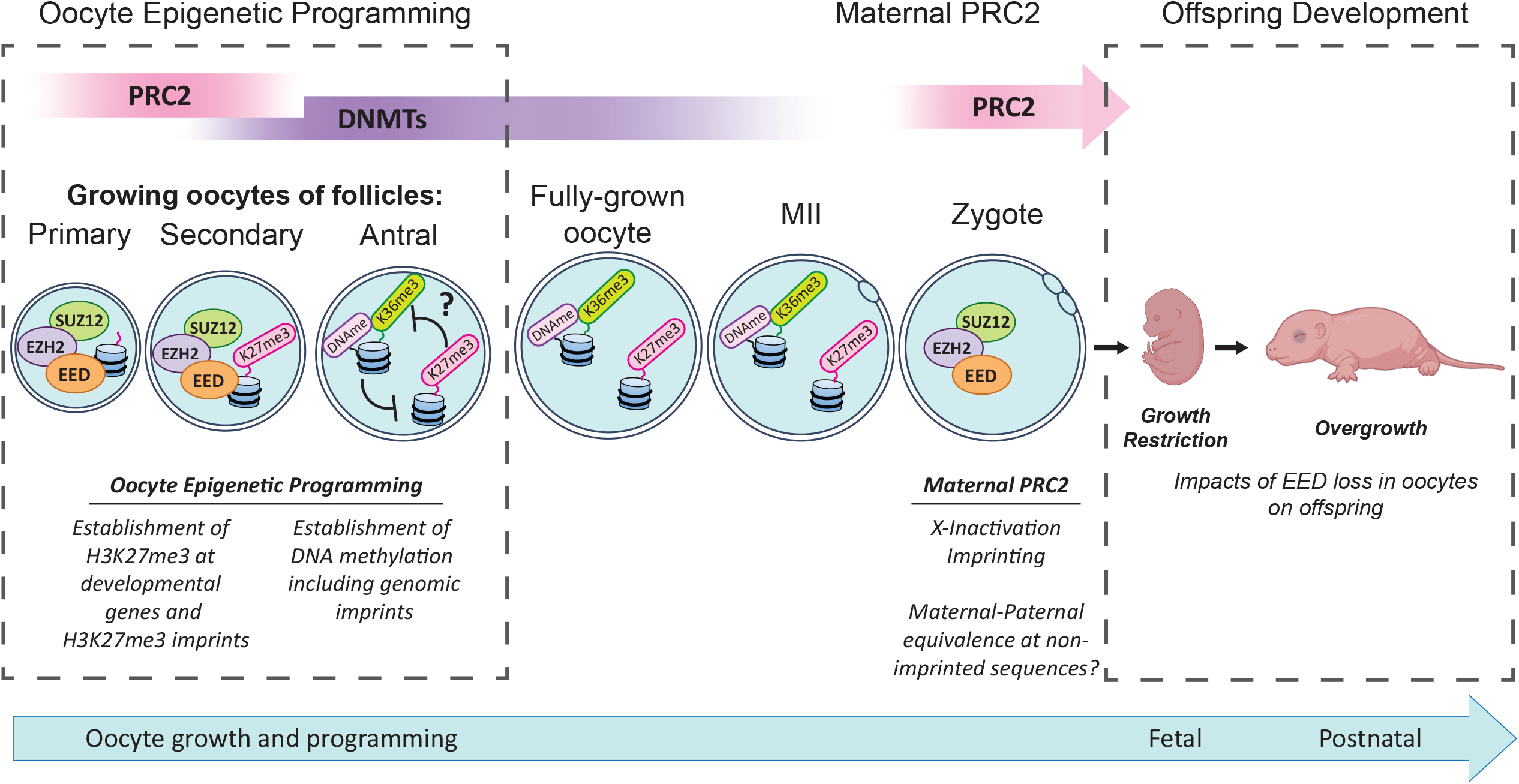
Summary of PRC2 functions during oocyte growth and maturation and pre-implantation development. All three essential components of PRC2 are present in growing oocytes only at the primary to secondary stages and establish H3K27me3 on H3K27me3 imprinted genes and a wide range of developmental genes, potentially programming *Eed* oocyte DEG expression in offspring. As this activity immediately precedes *de novo* DNA methylation, we propose that H3K27me3 established prior to DNA methylation may act as a “place-keeper” protecting developmental genes from modifications such as H3K36me3 and/or DNA methylation. Cytoplasmic PRC2 proteins and/or mRNA are inherited via the mature oocyte and regulate pre-implantation development, including X-inactivation, H3K27me3 dependent imprinting (31, 34, 35) and establishment of maternal – paternal equivalence at non-imprinted sequences. Loss of PRC2 in the oocyte leads to embryo growth restriction (33) but offspring are ultimately overgrown immediately after birth (40).

As transient PRC2 activity occurs early during oocyte growth and immediately precedes DNMT activity (61), it is reasonable to speculate that in the absence of H3K27me3 *Eed* oocyte DEGS may accumulate H3K36 and subsequent DNA methylation. Aberrant establishment of DNA methylation in *Eed* null oocytes may then contribute to developmental outcomes in subsequent offspring. To our knowledge, no study has directly measured the impact of maternal PRC2 deletion on global DNA methylation within the oocyte. Inoue *et al*. observed that classically imprinted gene expression was normal in maternal *Eed-*null embryos and concluded that DNA methylation establishment at classically imprinted genes was not impacted by *Eed* deletion (33). However, this does not exclude the possibility that H3K27me3 protects other regions from establishing DNA methylation in oocytes, particularly the oocyte DEGs identified in this study and other H3K27me3 enriched genes identified in other studies (33, 35, 44, 46). Further work is therefore required to determine the potential impact of H3K27me3 loss in primary-secondary oocytes on the broader epigenetic landscape of mature oocytes.

The potential for H3K27me3 to guide epigenetic state of target genes in oocytes is of interest, as using this model we previously observed post-natal overgrowth in offspring derived from oocytes lacking EED (40). However, another study showed that E10.5 embryos from *Eed*-null oocytes were growth restricted (33) indicating that loss of EED in oocytes differentially impacts embryonic and offspring growth. Placental hyperplasia has been attributed to loss of H3K27me3-dependent imprinting at a small number of genes in mice derived by SCNT (36-38), but is considered to be a placental effect. While this could lead to large offspring from oocytes lacking EED, it is yet to be observed in a model with oocyte-specific *Eed* deletion and does not explain why early embryos were smaller. One explanation could be that loss of maternal PRC2 in the zygote and early pre-implantation embryo hampers early development. However, an alternative explanation is that EED and H3K27me3-dependent programming of developmental genes in growing oocytes may also affect offspring growth and development through an as-yet undefined epigenetically inherited mechanism, such as altered DNA methylation.

Finally, previous studies have demonstrated a link between H3K27me3 and repression of repetitive sequences when DNA methylation levels are low, including in male and female fetal germ cells and embryonic stem cells (55-57). However, we did not observe any change in L1 expression in GV oocytes. With the obvious caveat that these sequences may have been repressed by other epigenetic modifications such as DNA methylation in growing oocytes, our data indicate that PRC2 is dispensable for repressing L1 sequences, and possibly other repetitive sequences in GV oocytes.

## Conclusions

In summary, we provide evidence that PRC2 acts transiently to establish H3K27me3 on a wide range of developmental genes in primary-secondary follicle oocytes and that this activity is required for the repression of these genes in fully-grown oocytes. As this activity precedes DNA methylation, and loss of H3K36me3 allows inappropriate spreading of H3K27me3 in oocytes, it seems likely that loss of H3K27me3 will affect other epigenetic programming events in oocytes. Moreover, as the transient activity of PRC2 in primary to secondary oocytes is distinct from the established maternal factor activity of PRC2 in pre-implantation embryos, and different gene sets are affected in oocytes and in pre-implantation embryos, these activities of PRC2 have distinct developmental consequences in offspring. Finally, as common genes are targeted for H3K27me3 enrichment in both mouse and human oocytes, understanding the activity of PRC2 during the growth of murine oocytes is likely to provide insight not only into non-genetic inheritance, but also for determining how altered PRC2 activity in oocytes affects human health.

## Methods

### Mouse strains, animal care and ethics

Mice were housed at Monash Medical Centre Animal Facility using a 12h light-dark cycle as previously reported (40). Food and water were available *ad libitum* with room temperature maintained at 21-23°C with controlled humidity. All animal work was undertaken in accordance with Monash University Animal Ethics Committee (AEC) approvals. Mice were obtained from the following sources: *Zp3Cre* mice (C57BL/6-Tg 93knw/J; Jackson Labs line 003651, constructed and shared by Prof Barbara Knowles (62)), *Eed* floxed mice (*Eed*^fl/fl^) (B6; 129S1-*Eed*tm1Sho/J; Jackson Labs line 0022727; constructed and shared by Prof Stuart Orkin *(63)*. The *Eed* line was backcrossed to a pure C57BL6/J and shared with us by Associate Professor Rhys Allen and Professor Marnie Blewitt, Walter and Eliza Hall Institute for Medical Research, Melbourne.

### Genotyping

All animals were genotyped via ear punch at weaning by Transnetyx (Cordova, TN) using real-time PCR assays (details available upon request) designed for each gene as described previously (40).

### Collection, antibody incubation and detection of ovaries for immunofluorescence

Ovaries for immunofluorescence (IF) were fixed in 4% paraformaldehyde (PFA) overnight at 4°C. Samples were then washed in PBS and processed through 70% ethanol and embedded in paraffin blocks, sectioned at 5μm and transferred to Superfrost™ Plus slides (Thermo-Fisher). Antigen retrieval was performed using DAKO Citrate buffer (pH 6.0) at 98°C for 30 minutes and non-specific binding blocked in PBS containing 5% BSA and 10% donkey serum for 1h at RT. Blocking solution was replaced with PBS containing 0.1% Triton-X 100 (PBSTX) and appropriately diluted primary antibodies (Supp. Materials and Methods) and incubated overnight at 4°C. Slides were washed in PBS and incubated with PBSTX containing secondary antibody for 1h at RT. After final washes in PBS, slides were rinsed in distilled H2O, mounted in DAPI ProLong Gold® and dried overnight. Fluorescence was detected using the VS120 Slide Scanner and quantified using QuPath Image Analysis Software (QuPath). Background fluorescence in the oocyte cytoplasm was removed from nuclear intensity. When comparing control versus experimental groups, the control mean was set to 1.0.

### Collection of oocytes and pre-implantation embryos for immunofluorescence

Ovaries were punctured using 30 G needles to release oocytes. GV oocytes were partially denuded mechanically using a narrow-bore glass pipette. For MII oocyte and zygote collections, females were injected with 5 international units (IU) Pregnant Mare Serum Gonadotropin (PMSG) followed by Human Chorionic Gonadotropin (hCG; 48h interval) and in the case of zygote collections were bred to C57BL/6 males. MII oocytes or zygotes were removed from the ampulla, denuded in M2 media containing hyaluronidase (0.3 mg/ml). Samples were either frozen and used for RNA analysis or fixed in 4% PFA containing 2% Triton X-100 for 30 minutes at RT. Samples were then washed in PBS containing 0.1% Tween, 0.01% Triton X-100 and 1% BSA (PBST-BSA) and stored in PBST-BSA.

### Oocyte and zygote whole-mount immunofluorescence

GV, MII oocytes and zygotes were blocked in PBST-BSA containing 10% donkey serum for 1 hour (h) at RT. The solution was then replaced with PBST-BSA containing appropriately diluted primary antibodies (Supp. Materials and Methods) and incubated overnight (o/n) at 4°C. Samples were washed in PBST-BSA and then incubated in PBST-BSA containing secondary antibodies (Supp. Materials and Methods) for 4h (GVs) or 1h (MII and zygote) in the dark at RT. After washing with PBST-BSA, samples were incubated with Hoechst 33342 (500 μg/ml) or DAPI (100 μg/ml) for 1h at RT, washed and stored in PBST-BSA until imaging. Fluorescence was detected using the Nikon C1 inverted confocal microscope and signal intensity quantified using ImageJ. Background fluorescence levels were measured in the cytoplasm and removed from nuclear intensity, with control mean was set to 1.0 for comparisons.

### Collection of oocyte RNA and RNA-sequencing

Cumulus-Oocyte Complexes (COCs) were collected from eight to twelve-week-old female mice and transferred to M2 media. Oocytes were denuded mechanically with a narrow-bore glass pipette and incubated with M2 media containing 5 μg/mL Hoechst 33342 for 10 minutes at 37°C.

GV oocytes were then scored as either surrounded nucleolus (SN), or non-surrounded nucleolus (NSN) based on Hoechst staining. SN oocytes were then collected, frozen on dry ice and stored at -80°C until RNA extraction. Ten to fifteen oocytes isolated from each female were pooled and total RNA isolated using the Agencourt RNAdvance Cell V2 extraction kit. High quality RNA (RIN >7.5) was used for library preparation (>1.2ng total RNA) using the Nugen Trio Library protocol, MU01440V2; 2017. 75bp single end sequencing was carried out on 4-6 libraries / genotype using Illumina NextSeq500 High output mode and v2.5 chemistry (Illumina Protocol 15046563 v02, Mar 2016) to collect >25M reads per sample.

### RNA-sequencing data analyses

Adaptor and low quality sequences in raw sequencing reads were trimmed either using AdaptorRemovel (64) (v2.2.1) for the oocyte RNA-Seq dataset with the following parameters: --trimns --trimqualities --minquality 20 --minlength 35, or using Trimmomatic (65) (v0.39) for bone growth plate and placental RNA-Seq datasets with the following parameters: LEADING:3 TRAILING:3 SLIDINGWINDOW:4:15 MINLEN:20. Clean reads were mapped to the mouse reference genome (GRCm38) using either STAR (66) (v2.5.3a) for the oocyte RNA-Seq dataset with default settings, or STAR (v2.7.5c) for the bone growth plate and placental RNA-Seq datasets with following settings: outFilterMismatchNoverLmax 0.03 --alignIntronMax 10000. For oocyte RNA-Seq dataset, raw counts for mouse reference genes (ensembl-release-93) were calculated using featureCounts (67) (v1.5.2) based on mapped bam files with the following parameters: -Q 10 –s 2. For bone growth plate and placental RNA-Seq datasets, raw counts for mouse reference genes (ensembl-release-101) were calculated using STAR (v2.7.5c) with parameter “--quantMode GeneCounts”. Raw counts of technical replicates from different sequencing lanes for the same samples in bone growth plate RNA-Seq dataset were merged. Differential gene expression analysis was carried out using R package “limma” (68) with “treat” function with parameter as “lfc=log(1.2)” for the oocyte RNA-Seq dataset or “lfc=log(1.1)” for the bone growth plate and placental RNA-Seq datasets. Statistically significantly differentially expressed genes were identified using “FDR < 0.05”. Gene Ontology (GO) enrichment analysis for significantly differentially expressed genes was carried out using The Database for Annotation, Visualisation and Integrated Discovery (DAVID) with following settings: GO term level 3, minimum gene count 5, and FDR < 0.05 (69).

### Analyses of genome-wide H3K27me3 distribution and H3K27me3 datasets in oocytes

*Eed* GV DEGs were compared to publicly available H3K27me3 ChIP-seq, CUT & RUN and CUT & TAG datasets of oocytes, sperm and pre-implantation embryos. Datasets used are summarised in Supp. Materials and Methods.

For the dataset from Zheng *et al*. 2016 (GSE76687), processed files including whole genome scale broad H3K27me3 peaks were downloaded and used for the comparison (44). For the dataset from Liu *et al*. 2016 (GSE73952), H3K27me3 states of the promoter regions of mouse reference genes were retrieved from Table S1 of the paper and used for the comparison (46). For the dataset from Erkek *et al*. 2013 (GSE42629), raw sequencing data were downloaded and then adaptor and low-quality sequences were trimmed using bbduk (v38.94) (47). Clean reads were mapped to the mouse reference genome (mm9) using bowtie2(70) (v2.4.4) with default settings. H3K27me3 peaks were identified using MACS2(71) (v2.1.1) with the following parameters: -g 1.87e9 --nomodel --broad -q 0.05. For the dataset from Inoue *et al*. 2018 (GSE116713), bigwig format files were downloaded and converted to bedGraph format using “bigWigToBedGraph” from UCSC utilities, and then H3K27me3 peaks were called using MACS2 (v2.1.1) based on the bedGraph files with the following parameter: -c 1.3 (equivalent to P value < 0.05) (33). For the human dataset from Xia *et al*. 2019 (GSE124718), processed files including whole genome scale broad H3K27me3 peaks were downloaded and used for the comparison (48).

This comparison took the 349 *Eed* GV DEGs and asked whether their promoters (defined as 2Kb up- or down-stream of TSS to be consistent with Liu *et al*. 2016) overlapped with H3K27me3 peaks (> 200bp overlap) in the above mentioned publicly available H3K27me3 ChIP or CUT & RUN datasets. For the human dataset from Xia *et al*. 2019 (GSE124718), human orthologous genes of mouse 349 *Eed* GV DEGs were identified and used for the comparison.

### Statistical Analyses

GraphPad Prism 9 was used for statistical analysis and to graph datasets. As appropriate, parametric Student’s *t* tests or ANOVA or non-parametric equivalents as indicated in figure legends.

## Supporting information

Jarred etal Supplementary Files

## Declarations

### Ethics approval and consent to participate

All animal work was undertaken in accordance with Monash University Animal Ethics Committee (AEC) approvals.

### Consent for publication

Not applicable

### Availability of data and materials

All RNA sequencing data have been deposited to the Gene Expression Omnibus (GEO) and are publicly available with accession number GSE193582. All other information is available from the corresponding author.

### Funding

This work was supported by grants and research funding from:

National Health and Medical Research Project Grant GNT1144966 (PSW, DKG, MvdB, DLA)

Hudson Institute of Medical Research (PSW)

Victorian Government’s Operational Infrastructure Support Program.

Monash University Postgraduate Student Awards (EGJ, RO and SP)

### Author contributions

Conceived and/or designed experiments/interpreted outcomes: EGJ, TT, ZQ, RO, SP, WTG, DLA, PSW

Performed experiments and/or analysed data: EGJ, ZQ, TT, RO, SP, HB, QZ, JMS, DLA, PSW

Bioinformatic analyses and/or comparisons with published datasets: ZQ, EGJ, SMP, DLA, PSW

Resources and/or supervision PSW DKG, MvdB, DLA, JC, NAS

Writing - original draft: EGJ, PSW, ZQ, RO, SP, NAS, DLA

Writing - review & editing: EGJ, PSW, RO, SP, DLA. All authors critically read and approved the manuscript.

### Data and materials availability

With exception of the RNA sequencing data generated in this study, all data are available in the main text, the Supplementary Materials. The RNA sequencing data have been deposited in the Gene Expression Omnibus (GEO) and are publicly available with accession number GSE193582. Published datasets used in this study are summarised in the Supplementary materials.

## Acknowledgments

We thank Dr. Neil Youngson and Prof. Marnie Blewitt for critical comments on the manuscript, Monash Animal Research Platform staff for assistance with mouse care, Monash Histology Platform for assistance with slide scanning and the Monash Micro Imaging Facility and MHTP Medical Genomics Facilities for assistance and technical advice.

