## Supplementary material for "Transient Polycomb activity represses developmental genes in growing oocytes": Jarred etal Supplementary Files

**Affiliations:** [1]Centre for Reproductive Health, Hudson Institute of Medical Research and Department of Molecular and Translational Science, Monash University, Clayton, Victoria, Australia; [2]Department of Bioinformatics and Computational Genetics, School of Biological Sciences, University of Adelaide, Adelaide, South Australia, Australia; [3]Biomedicine Discovery Institute, Monash University, Clayton, Victoria, Australia; [4]School of BioSciences, University of Melbourne, Parkville, Victoria, Australia; [5]School of Psychology and Public Health, La Trobe University, Melbourne, Victoria, Australia; [6]Bone Cell Biology and Disease Unit, St. Vincent's Institute of Medical Research and Department of Medicine at St. Vincent's Hospital, Fitzroy, Victoria, Australia; [7]Department of Medical Genetics, University of British Columbia and British Columbia Children's Hospital Research Institute, Vancouver, BC, Canada.

References are included in the main manuscript

**Please note: Supplementary Tables 1-10** are included as a separate Supplementary excel file with this submission

**Supplementary Figures:**

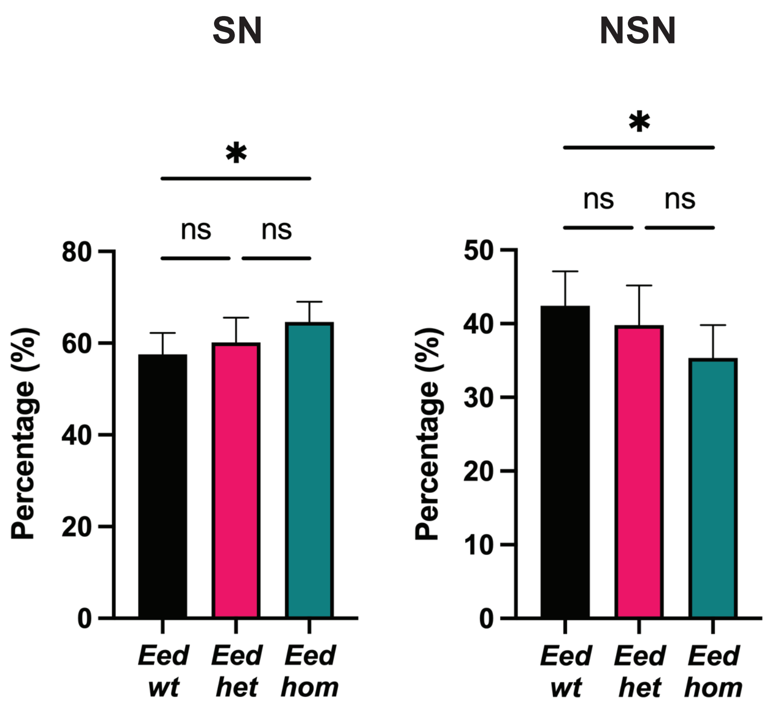
SN
NSN
*
ns
ns
*
ns
ns
80
60
40
20
0
50
40
30
20
10
0
Percentage (%)
Percentage (%)
Eed wt
Eed het
Eed hom
Eed wt
Eed het
Eed hom

Supp. Fig. 1

**Supp. Fig. 1. Deletion of *Eed* in oocytes moderately increased the rate of Surrounded Nucleolus (SN) GV oocytes compared to Non-Surrounded Nucleolus (NSN) GV oocytes.** Percentage of SN (left) and NSN (right) GV oocytes obtained from *Eed-wt, Eed-het,* and *Eed-hom* females during oocyte collections. *P < 0.05, one-way ANOVA plus Tukey's multiple comparisons test, N = 7 *Eed-wt*, 5 *Eed-het* and 7 *Eed-hom* females. Error bars represent mean ± standard deviation.

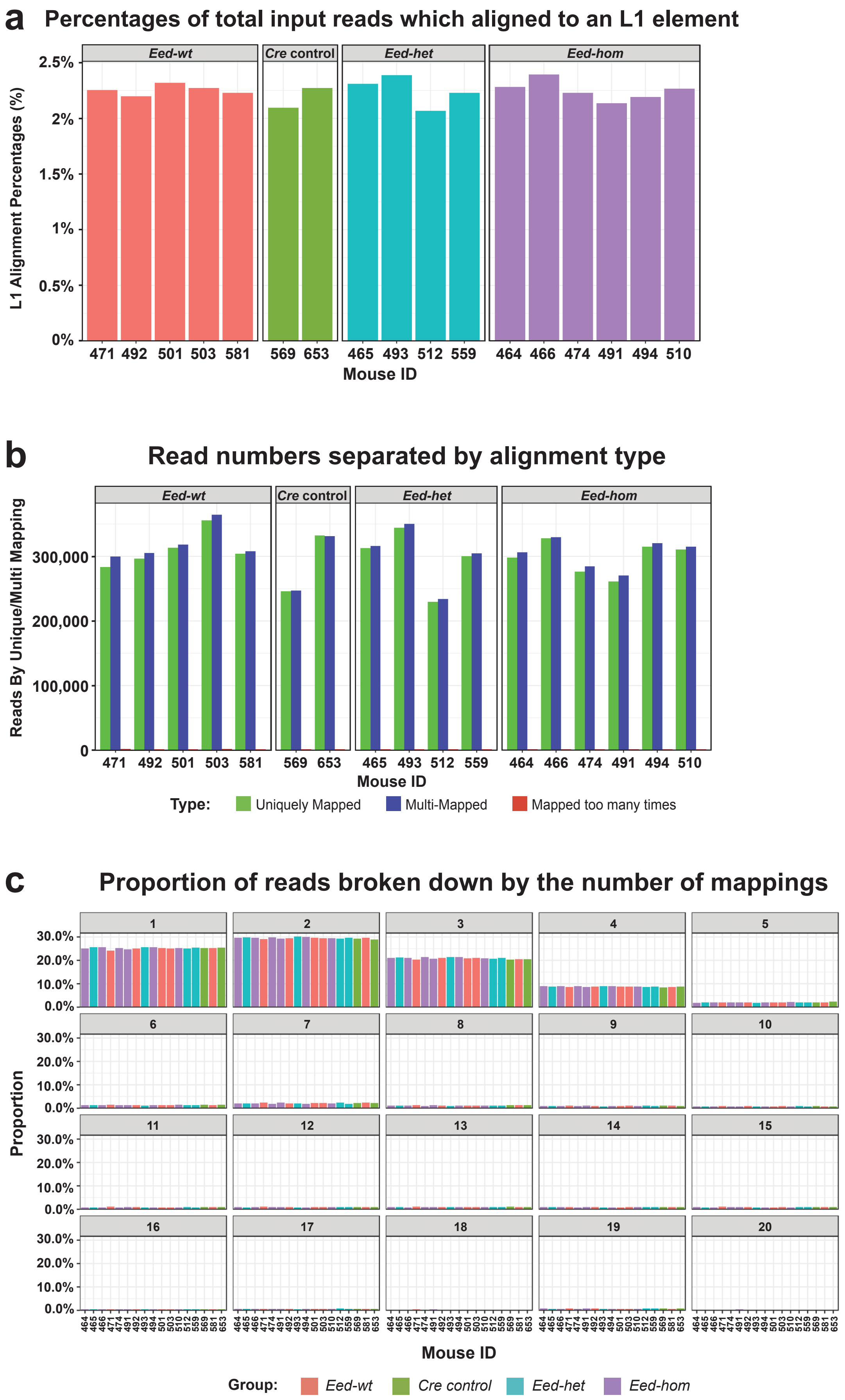
a
Percentages of total input reads which aligned to an L1 element
Eed-wt
Cre control
Eed-het
Eed-hom
L1 Alignment Percentages (%)
2.5%
2%
1.5%
1%
0.5%
0%
471 492 501 503 581
569 653
465 493 512 559
464 466 474 491 494 510
Mouse ID
b
Read numbers separated by alignment type
Reads By Unique/Multi Mapping
300,000
200,000
100,000
0
Mouse ID
Type: Uniquely Mapped Multi-Mapped Mapped too many times
c
Proportion of reads broken down by the number of mappings
Proportion
30.0%
20.0%
10.0%
0.0%
Mouse ID
Group: Eed-wt Cre control Eed-het Eed-hom

Supp. Fig. 2

**Supp. Fig. 2. Loss of *Eed* in growing oocytes did not impact expression of LINE-1 transposons.** **(a)** Percentages of total input reads which aligned to a LINE-1 (L1) element. **(b)** Number of reads which map to unique and multiple L1s. **(c)** Proportions of reads according to the number of sites mapped to per read, to a maximum of 20. For **(a-c)**, data represent individual replicates from *Eed-wt* (n = 5), *Eed-wt Cre* (n = 2), *Eed-het* (n = 4) and *Eed-hom* (n = 6) females.

**_Eed_ DEGs vs pre-implantation DEGs**

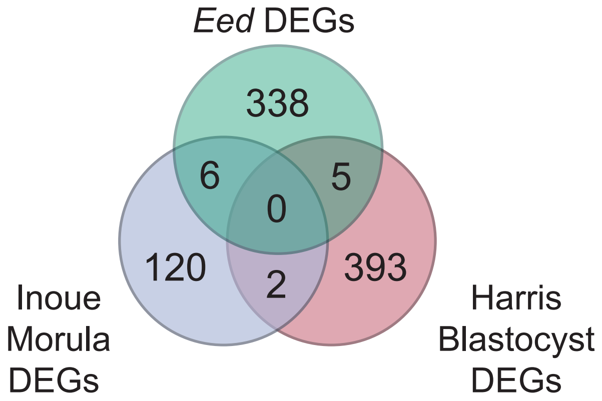


**Supp. Fig. 3**

**Supp. Fig. 3. *Eed* oocyte DEGs were not dysregulated in pre-implantation embryos**. Venn Diagram comparing *Eed* oocyte DEGs against DEGs identified in *Eed* maternal null morula and blastocyst embryos. Six genes (*Plxnd1*, *Tceal8*, *Rap2c*, *Bbx*, *Xlr3c* and *Trm2b*) were common in oocytes and morula embryos, five genes (*Chrdl1*, *Lonrf2*, *Trim6*, *Cyp1b1* and *Ccbe1*) were common in oocytes and blastocyst embryos, and two genes (*Tspan6* and *Gk*) were common in morula and blastocyst embryos. For full DEG lists see Tables S1, S6 and S7. Morula and Blastocyst datasets were generated by analysis of published raw datasets (Manuscript References 33, 34).
